# Characterization of the Drosophila adult hematopoietic system reveals a rare cell population with differentiation and proliferation potential

**DOI:** 10.1101/2021.07.06.451161

**Authors:** Manon Boulet, Yoan Renaud, François Lapraz, Billel Benmimoun, Laurence Vandel, Lucas Waltzer

## Abstract

While many studies have described Drosophila embryonic and larval blood cells, the hematopoietic system of the imago remains poorly characterized and conflicting data have been published concerning adult hematopoiesis. Using a combination of blood cell markers, we show that the adult hematopoietic system is essentially composed of a few distinct mature blood cell types. In addition, our transcriptomics results indicate that adult and larval blood cells have both common and specific features and it appears that adult hemocytes reactivate many gene expressed in embryonic blood cells. Interestingly, we identify a small set of blood cells that do not express differentiation markers but maintain the progenitor marker *domeMeso*. Yet, we show that these cells are derived from the posterior signaling center, a specialized population of cells present in the larval lymph gland, rather than from larval blood cell progenitors, and that their maintenance depends on the EBF transcription factor Collier. Furthermore, while these cells are normally quiescent, we find that some of them can differentiate and proliferate in response to bacterial infection. In sum, our results indicate that adult flies harbor a small population of specialized cells with limited hematopoietic potential and further support the idea that no substantial hematopoiesis takes place during adulthood.

## Introduction

Several aspects of blood cell development and functions have been conserved between mammals and Drosophila (Banerjee et al., 2019; Hartenstein, 2006). Hence this insect has been used as a simple genetic model organism to study the fundamental bases underlying hematopoiesis and blood cell functions (Boulet et al., 2018; Letourneau et al., 2016). Yet, while much effort has been devoted to the characterization of the Drosophila blood cells at the embryonic and larval stages, the adult stage remains less well characterized and conflicting data have been published (see below). Thus, further work is essential to gain a better understanding of the composition and dynamic of the adult hematopoietic system.

As in vertebrates, Drosophila hematopoiesis occurs in successive waves and stems from mesoderm-derived blood cell progenitors (prohemocytes) (Banerjee et al., 2019). These blood cell progenitors can give rise to three main differentiated cell types, collectively called hemocytes and related to the vertebrate myeloid cells: plasmatocytes, crystal cells and lamellocytes (Gold and Bruckner, 2015; Parsons and Foley, 2016). Plasmatocytes are macrophages that form the bulk of the population; they are essentially implicated in tissue remodeling and in the cellular immune response, while crystal cells are implicated in clotting and melanization (an insect-specific immune response). Lamellocytes are normally barely present but their differentiation is massively induced in the larvae in response to pathological situations, such as infestation by parasitoid wasp eggs.

Plasmatocytes and crystal cells are first produced in the embryo after a small group of prohemocytes emerge from the early head mesoderm and differentiate (Bataille et al., 2005; Lebestky et al., 2000; Tepass et al., 1994). These embryo-derived hemocytes compose the larval peripheral blood cells, which are observed as free circulatory cells in the hemolymph or attached under the cuticle (Holz et al., 2003; Makhijani et al., 2011). Interestingly, differentiated larval plasmatocytes can not only proliferate but also transdifferentiate into crystal cells (Leitao and Sucena, 2015; Makhijani et al., 2011), and it was proposed that the increase in peripheral blood cells during larval development relies on self-renewing plasmatocytes (Gold and Bruckner, 2015). Yet, undifferentiated blood cells were also described among peripheral hemocytes (Sinenko et al., 2010) and recent single-cell sequencing experiments suggest that proliferative plasmatocytes retain a progenitor signature (Cattenoz et al., 2021; Cattenoz et al., 2020; Tattikota et al., 2020). In parallel, a second wave of hematopoiesis takes place during the larval stages in a specialized organ called the lymph gland (Lanot et al., 2001). The lymph gland precursors derive from the embryonic lateral mesoderm and develop in close association with the anterior part of the dorsal vessel (Crozatier et al., 2004; Mandal et al., 2004). In third instar larvae, the lymph gland is composed of 3 to 4 pairs of lobes separated by pericardial cells (Lanot et al., 2001; Rodrigues et al., 2021). The posterior lobes are essentially composed of prohemocytes (Rodrigues et al., 2021), while the anterior lobes contain blood cells progenitors, their differentiated progenies (plasmatocytes and crystal cells), and a small cluster of cells that form a niche, called the posterior signaling center (PSC) (Jung et al., 2005; Letourneau et al., 2016). In normal situations, lymph gland blood cells are released into circulation during pupation when the lymph gland disperses (Grigorian et al., 2011; Honti et al., 2010).

By comparison, the adult hematopoietic system of the fly remains poorly characterized (Banerjee et al., 2019). Initial examinations failed to reveal the presence of a hematopoietic organ but showed that hemocytes are scattered throughout the imago, mostly as sessile populations, and accumulate along the heart in the abdomen (Elrod-Erickson et al., 2000; Lanot et al., 2001), a general feature among insects (Yan and Hillyer, 2020). In addition, it seemed that all the hemocytes were non-dividing phagocytic cells whose number and phagocytic activity decrease with age (Lanot et al., 2001; Mackenzie et al., 2011; Woodcock et al., 2015). Besides, transplantation experiments showed that both the embryonic and the larval hematopoietic anlagen contribute to the adult blood cells (Holz et al., 2003). Hence, the prevailing view was that the adult hematopoietic system was solely composed of mature plasmatocytes derived from the embryonic and larval stages, and that no hematopoiesis took place during adulthood. Yet, some data suggested a more contrasted situation. First, the analysis of different markers revealed that crystal cells and distinct subpopulation of plasmatocytes are present in the adult (Clark et al., 2011; Kurucz et al., 2007). Second, a seminal study by Ghosh *et al*. suggested that hematopoiesis also occurs in the imago. Indeed, the authors proposed that blood cell progenitors capable of differentiation persist in the adult and they observed plasmatocyte proliferation in response to *E. coli* infection (Ghosh et al., 2015). However, these findings were rebuked by a subsequent publication (Sanchez Bosch et al., 2019), in which the authors refuted the claim that the GATA factor Serpent (Srp) is a marker of adult blood cell progenitors and found no evidence for blood cell proliferation or differentiation.

Here, we set out to better characterize the Drosophila adult hematopoietic system. First, we established the gene expression profile of adult hemocytes and compared it to their larval ascendant (larval peripheral hemocytes and lymph gland) to define the common and specific features of these immune cells. Next, we used a panel of hematopoietic markers to assess their expression in the imago and gain a better appreciation of the adult blood cell landscape diversity and evolution with age. Finally, we focused our analysis on a small population of adult blood cells that do not express hemocyte differentiation markers. We show that they originate from the PSC rather than from larval blood cell progenitors and that they can proliferate and differentiate in response to infection albeit unfrequently. These PSC-derived cells most probably account for the previously reported population of adult blood cell progenitor. Our findings are discussed in view of the current controversies in the field.

## Results

### Adult blood cells exhibit a distinct gene expression profile as compared to larval blood cells

As a first step to characterize Drosophila adult hematopoietic system, we used a perfusion protocol similar to the one used to bleed mosquitoes (Muller et al., 1999) in order to retrieve adult blood cells and define their gene expression program (see Materials and Methods). We found that ±99% of the cells collected with this technique in 5-day old flies expressed the panhemocyte marker *srp* as revealed using the *srpD-GAL4* driver (Waltzer et al., 2002) or the *srpHemo-H2A::3mCherry* reporter (Gyoergy et al., 2018) (Figure 1A, C, E), indicating that this approach allows to obtain clean blood cell preparations. Of note, when we used young adults (24-48 hours after hatching), the bleeds were contaminated with remnants of the larval fat body (Figure 1B, arrow), which is known to dissociate into large individual adipose cells during metamorphosis (Nelliot et al., 2006) and gets cleared in the first two days of adulthood (Aguila et al., 2007). Using aged flies (5-week old), we observed that ±4,6% of the cells did not express *srp* and we recovered fewer hemocytes (Figure 1D-F), which is consistent with previous reports showing that blood cell number decreases with aging (Mackenzie et al., 2011; Sanchez Bosch et al., 2019; Woodcock et al., 2015).

**Figure 1.**
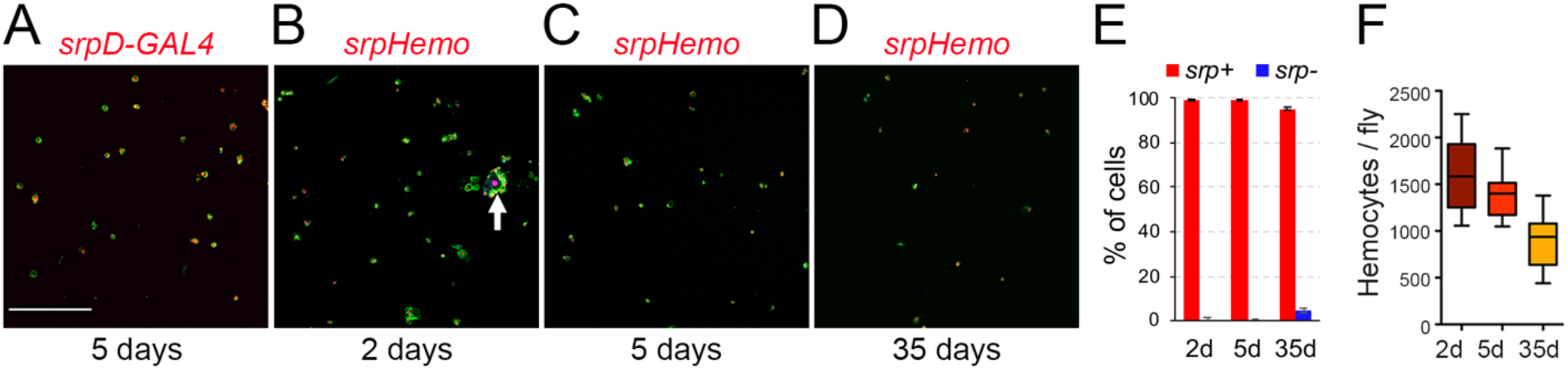
Recovery of Drosophila blood cell by bleeding. **(A-D)** Confocal images of cells recovered from *srpD-GAL4,UAS-mCherry* (A) or *srpHemo-His2A-RFP* (B-D) adult females. Cells were counterstained with phalloidin (green) and DAPI (blue). The age of the flies is indicated below each panel. Scale bar: 200µm. The arrow in B indicates a fat body cell surrounded by hemocytes. **(E)** Quantifications of the proportion of cells (DAPI^+^) expressing *srpHemo-His2A-RFP* according to fly age. **(F)** Quantifications of the absolute number of cells recovered per fly at different ages.

Next, we established the gene expression profile of adult hemocytes collected from 5-day old flies by RNA sequencing. For comparison, we also established the transcriptome of 3^rd^ instar larval peripheral hemocytes (*i.e.* embryo-derived hemocytes) and lymph gland hemocytes. All experiments were performed with biological triplicates and we obtained at least 14 million mapped reads per sample. We found that 6802 genes are expressed with a RPKM>1 in all three samples of adult hemocytes (corresponding to 39% of the genes on Drosophila reference genome dm6) (Supplementary Table 1). Similarly, larval peripheral blood cells and lymph gland hemocytes expressed respectively 6282 and 6220 genes (Supplementary Table 2 and 3). As shown in Figure 2A, adult hemocytes, lymph gland and larval peripheral hemocytes shared the expression of 5351 genes, *i.e.* between 79 and 86% of their respective transcriptome. Among the 268 genes annotated in Flybase (FB_2021_02) as being expressed in blood cells (*i.e.* hemocytes, plasmatocytes, crystal cells, lamellocytes or prohemocytes, Supplementary Table 4), 216 were retrieved in our adult hemocyte data set (±2.1-fold enrichment, p-value<5.5E-45), 197 in the lymph gland (±2.1-fold enrichment, p-value<3.5E-37), and 198 in peripheral hemocytes (±2.0-fold enrichment, p-value<3.4E-37), with 188 being common to all three samples (Supplementary Figure 1A). Thus, most hemocyte markers expressed in larval blood cells are also expressed in the adult. Notably, 11 of these hemocyte markers (*Col4a1/Cg25C, Ppn, Nplp-2, Pxn, Fer2LHC, Fer1HCH, PPO2, PPO1, crq, Sr-Cl* and *Glt)* were among the 60-most expressed genes in adult blood cells (Supplementary Table 1). In contrast, *gcm* and *gcm2,* which were shown to be expressed in embryonic but not in larval blood cells (Avet-Rochex et al., 2010; Bazzi et al., 2018; Ramond et al., 2020), were not detected in adult (or larval) hemocyte RNA-seq samples (Supplementary Tables 1-3).

**Figure 2.**
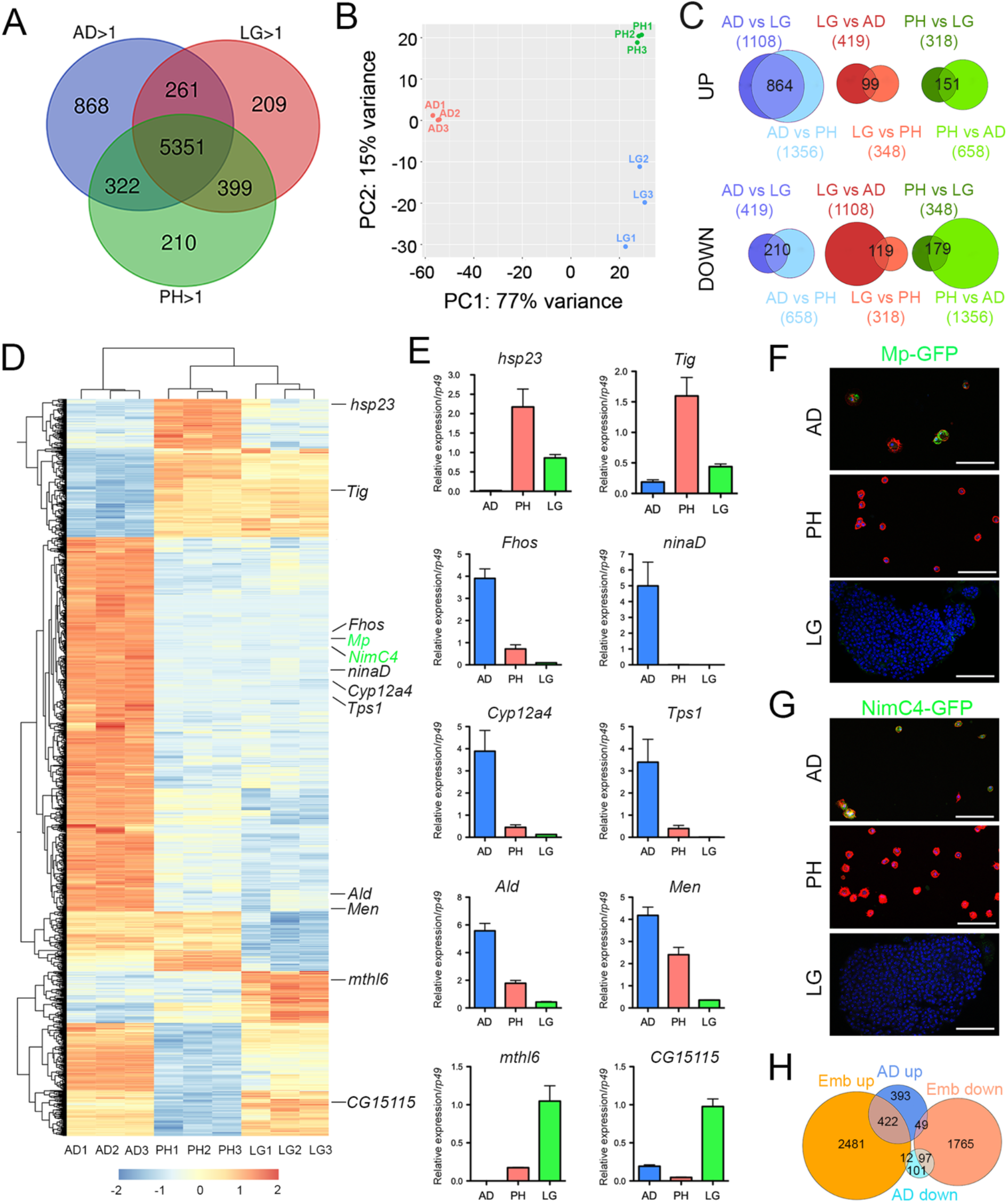
Analysis of adult blood cells gene expression profile. **(A)** Venn diagrams showing the number of genes expressed (RPKM>1 in all three biological replicates) in adult hemocytes (AD), larval peripheral hemocytes (PH) and larval lymph glands (LG) as determined by RNA-seq on *w^1118^* females. **(B)** Principal component analysis of the RNA-seq profiles of adult, peripheral and lymph gland biological triplicates. **(C)** Venn diagrams showing the overlaps between differentially expressed genes in pairwise comparisons (adjusted p-value<0,01 and fold change >2). Upper panels: up-regulated genes. Lower panels: down-regulated genes. The number of up or down-regulated genes in each condition is indicated between brackets and the number of genes commonly deregulated is indicated in the Venn diagram intersection. **(D)** Heat map of the genes differentially expressed between adult *versus* larval hemocytes. The genes tested in panels E to G are indicated. **(E)** The expression level of the indicated genes was determined by RT-qPCR on adult hemocytes, larval peripheral hemocytes and larval lymph glands. Expression levels were normalized to *rp49*. Means and SEM from 3 independent experiments are represented. **(F-G)** Confocal images showing GFP expression in adult and larval hemocytes from *Mp-GFP* (F) or *NimC4-GFP* (G) reporter lines. Nuclei were stained with DAPI (blue). For adult and larval peripheral hemocytes (upper and middle panels), the samples were stained with phalloidin (red). Lower panels show one LG anterior lobe. Scale bar: 50µm. **(H)** Venn diagrams showing the overlaps of up or down-regulated genes between adult or embryonic (Emb) hemocytes *versus* larval hemocytes.

Although they express many genes in common, adult and larval hemocytes also exhibit distinct features. Indeed, principal component analysis showed that 77% of gene expression variance among samples was accounted by stage differences (adult *versus* larva) (Figure 2B). Furthermore, differential gene expression analysis with DESeq2 revealed that 1074 genes are differentially expressed between adult blood cells and both larval peripheral hemocytes and lymph glands (adjusted p-value<0.01 and fold change >2), with 864 genes up-regulated and 210 down-regulated in adult hemocytes (Figure 2C, D and Supplementary Table 5). For comparison, 330 genes were differentially expressed between larval peripheral hemocytes and both adult blood cells and lymph glands (151 up-regulated, 179 down-regulated), and 218 genes were differentially expressed between lymph glands and both adult and larval peripheral hemocytes (99 up-regulated, 119 down-regulated). For instance, consistent with previous immunostaining results (Kurucz et al., 2007), we observed that *hemese* expression was down-regulated in adult hemocytes as compared to larval ones (Supplementary Table 5). In addition, to validate these results, we performed RT-qPCR on a few genes that were differentially expressed between adult and larval blood cells using independent RNA samples. Thereby we confirmed that *hsp23, Tig, mthl6* and *CG15115* were downregulated in adult hemocytes whereas *ninaD, Fhos, Tps1, Cyp12a4*, *Ald* and *Men* were up-regulated (Figure 2E). Furthermore, reporter lines for *NimC4* and *Mp* also showed that these two genes are expressed in adult hemocytes but not in larval ones (Figure 2F,G).

Gene ontology enrichment analyses underlined the up-regulation of genes implicated in small molecule metabolism project, including ATP metabolism (as seen with the overexpression of several enzymes involved in glycolysis, TCA cycle, or oxidative phosphorylation) and a down-regulation of genes regulating chromatin organization in adult hemocytes (Table 1). Along the same line, Gene Ontology analysis considering the 1000 most strongly expressed genes in larval, lymph gland or adult blood cells revealed shared features between them, with for instance a strong over-representation for genes implicated in translation, vesicle-mediated transport or immune response, but also confirmed the singular enrichment for ATP metabolic processes among highly-expressed genes in adult hemocytes (Table 2).

**Table 1.** Main gene ontology terms enriched in genes over- (up AD) or under- (down AD) expressed in adult hemocytes as compared to larval ones. The p-value and the number of genes in each category are indicated.

| GO-term | up AD | down AD |
| --- | --- | --- |
| small molecule metabolic process | 5.06E-42 (n=158) |  |
| ATP metabolic process | 1.66E-39 (n=63) |  |
| inner mitochondrial membrane protein | 3.50E-30 (n=52) |  |
| oxidoreductase activity | 7.74E-29 (n=113) |  |
| nucleosome |  | 6.27E-53 (n=45) |
| chromatin organization |  | 5.46E-29 (n=51) |

**Table 2.**
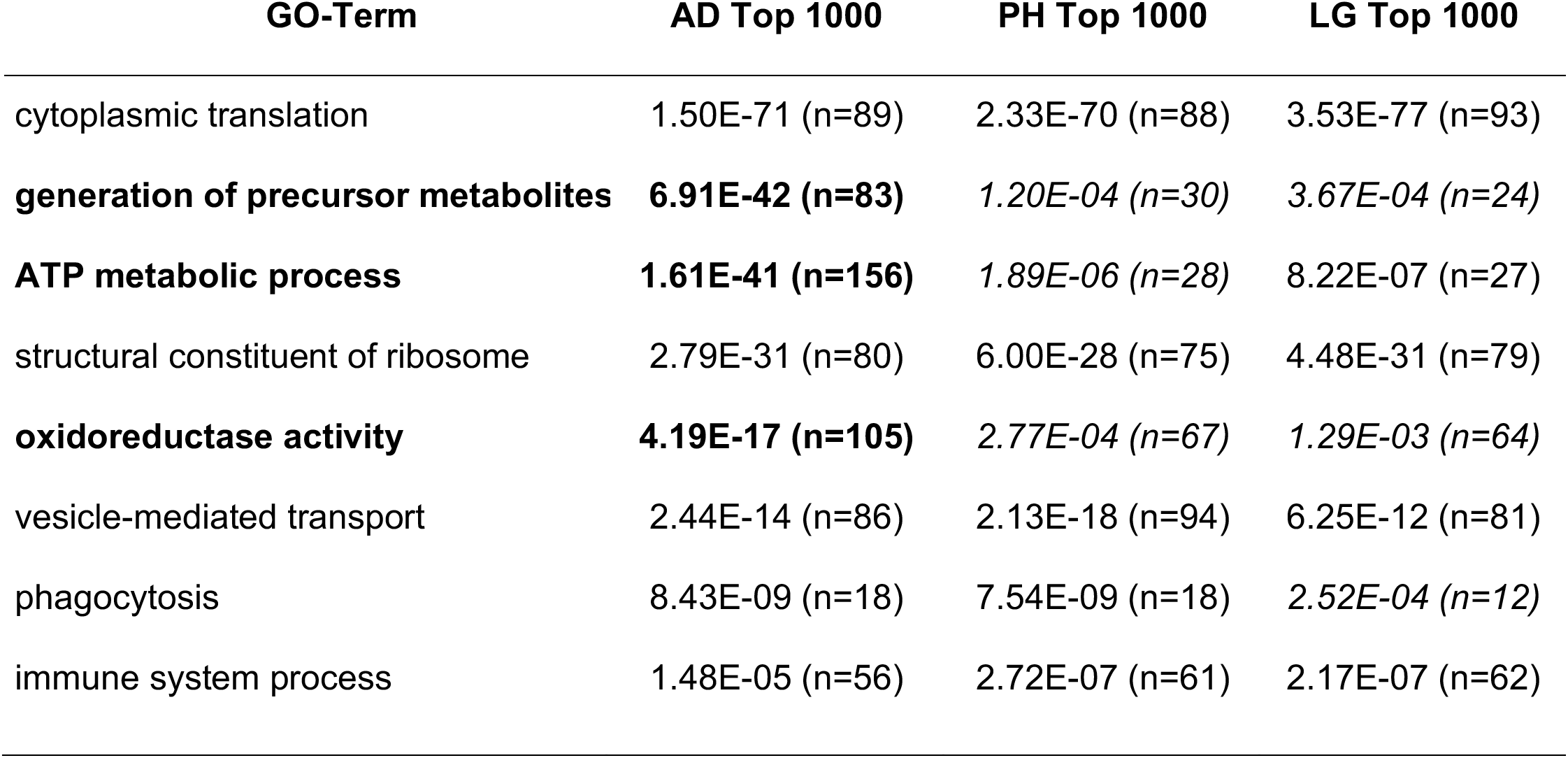
Main gene ontology terms among the 1000 most strongly expressed genes in adult hemocytes (AD), peripheral hemocytes (PH) and lymph glands (LG). The p-value and the number of genes in each category are indicated. Categories particularly enriched in adult hemocytes are highlighted in bold.

| GO-Term | AD Top 1000 | PH Top 1000 | LG Top 1000 |
| --- | --- | --- | --- |
| cytoplasmic translation | 1.50E-71 (n=89) | 2.33E-70 (n=88) | 3.53E-77 (n=93) |
| **generation of precursor metabolites** | **6.91E-42 (n=83)** | *1.20E-04 (n=30)* | *3.67E-04 (n=24)* |
| **ATP metabolic process** | **1.61E-41 (n=156)** | *1.89E-06 (n=28)* | 8.22E-07 (n=27) |
| structural constituent of ribosome | 2.79E-31 (n=80) | 6.00E-28 (n=75) | 4.48E-31 (n=79) |
| **oxidoreductase activity** | **4.19E-17 (n=105)** | *2.77E-04 (n=67)* | *1.29E-03 (n=64)* |
| vesicle-mediated transport | 2.44E-14 (n=86) | 2.13E-18 (n=94) | 6.25E-12 (n=81) |
| phagocytosis | 8.43E-09 (n=18) | 7.54E-09 (n=18) | *2.52E-04 (n=12)* |
| immune system process | 1.48E-05 (n=56) | 2.72E-07 (n=61) | 2.17E-07 (n=62) |

Surprisingly, one of the top-enriched gene in adult hemocytes is *NimC4*, which was recently shown to be highly enriched in stage 16 embryonic hemocytes as compared to larval descendants (*i.e.* larval peripheral hemocytes) in a comparative transcriptomic analysis (Cattenoz et al., 2020). Moreover, reminiscent of the situation that we observed, embryonic and their larval progenies were found to have distinct metabolic gene signature. We thus reanalyzed the RNA-seq data from (Cattenoz et al., 2020) using the same settings as for our transcriptomes. Thereby we established a list of 4892 differentially expressed genes between embryonic and peripheral hemocytes (adjusted p-value<0.01 and fold change >2; Supplementary Table 6) that we compared with our results. Strikingly, 49% (422/864) of the genes up-regulated in adult *versus* larval hemocytes are also over-expressed in embryonic blood cells (±2.9-fold over-enrichment, p<4.8E-113) (Figure 2H). Similarly, 46% (97/210) of the genes down-regulated in adult versus larval hemocytes are also repressed in embryonic blood cells (±4.2-fold over-enrichment, p<1.3E-38) (Figure 2H). The common over-expressed genes in embryonic and adult hemocytes *versus* larval ones included were enriched in genes implicated in small molecule/ATP metabolic processes, whereas down-regulated one were enriched for genes implicated in cell cycle or protein folding. Hence, many genes and process differentially regulated between adult and larval hemocytes are similarly regulated between embryonic and larval hemocytes.

In sum, these data show that adult hemocytes share a significant portion of their transcriptome with larval blood cells but also exhibit distinct features, some of which appear to be shared with embryonic hemocytes.

### Characterization of the adult hematopoietic system landscape

To better define the cellular composition of the adult hematopoietic system, we then analyzed the expression of a series of well-characterized reporters classically used to study embryonic and larval blood cell types (see Material and Methods).

First, we assessed the expression of plasmatocyte and crystal cell differentiation markers. In 5-days old flies, the plasmatocyte reporters for *Peroxidasin (Pxn)*, *Cg25C* (Collagen 25C, also known as *Collagen type IV alpha1), Hemolectin (Hml)* and *croquemort (crq)* were expressed in ±95%, 90%, 65% and 25% of the blood cells respectively, while those for the crystal cell markers *Black cells* (*Bc*, also known as *Prophenoloxydase 1, PPO1)* and *lozenge (lz)* were expressed in ±8 to 12% of the cells (Figure 3A-G and Supplementary Figure 1A-F). The proportion of hemocytes expressing these reporters was similar in 35-day old flies (Figure 3G). The analysis of double transgenic lines revealed that (almost) all the cells expressing *Hml* co-expressed *srp* and *Pxn* (Figure 3H, K and Q). Besides, ±90 of the *crq^+^* cells were *Hml^+^* (Figure 3O and Q). Among the *Bc^+^* cells, 100% expressed *srp* and *Pxn*, ±90% *Cg25C* and 60% *Hml* (Figure 3I, L, N, P and Q). Of note, it was previously observed that the “plasmatocyte” markers *Cg25C*, *Pxn* and *Hml* are also expressed in larval crystal cells (Jung et al., 2005). Again, we did not observe major shifts in the repartition of these populations between 5 and 35-day old flies, indicating that the overall composition of the adult hematopoietic system remains stable during aging (Figure 3Q).

**Figure 3.**
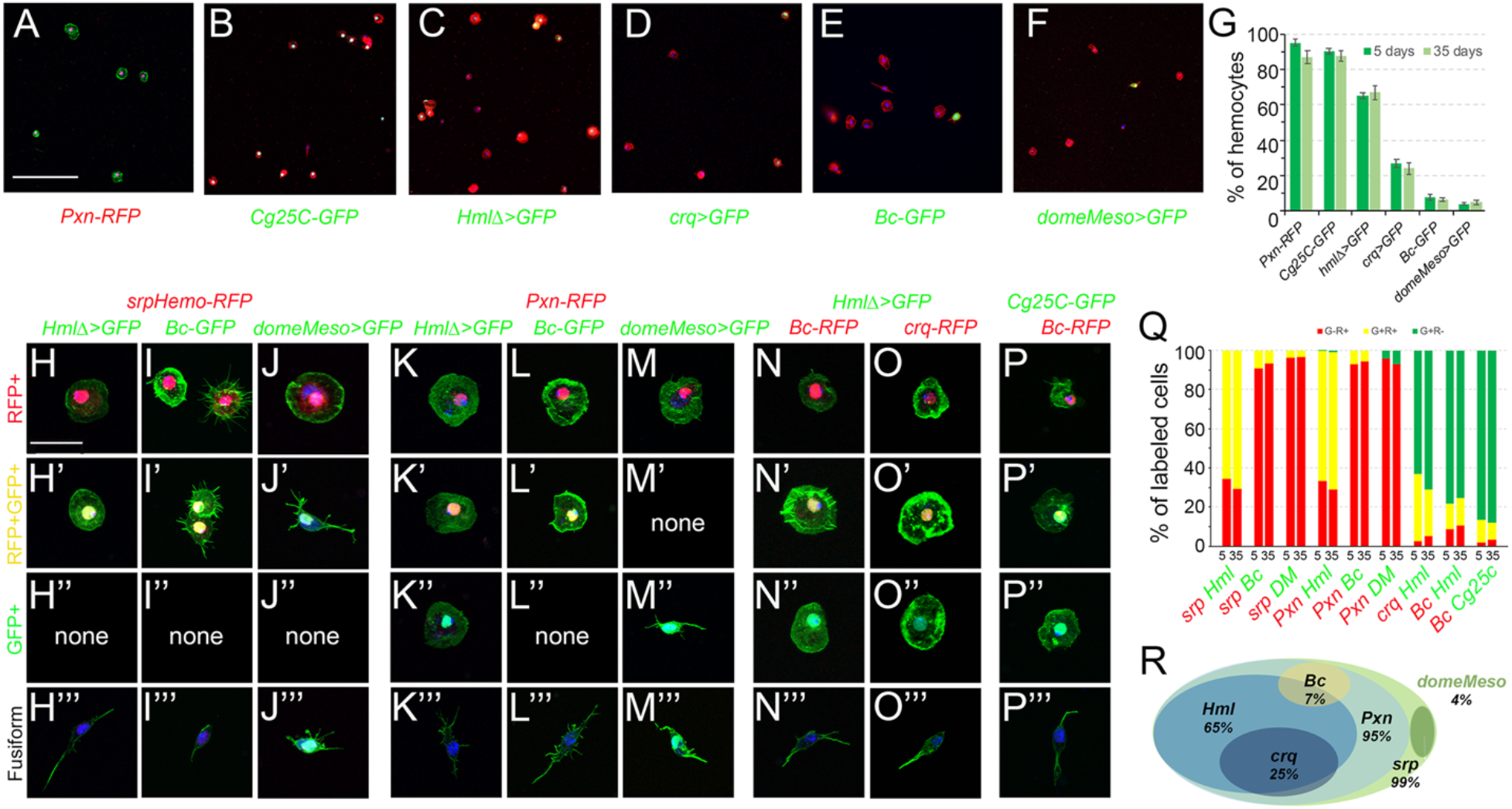
Analysis of blood cell transgenic markers in adult bleeds. **(A-F)** Confocal views showing representative bleeds from 5-day old flies carrying the following transgenes: *Pxn-RedStinger* (A, *Pxn-RFP*), *Cg25C-GFP* (B), *HmlΔ-GAL4,UAS-2xEYFP* (C, *HmlΔ>GFP*), *crq-GAL4,UAS-2xEYFP* (D, *crq>GFP*), *BcF2-GFP* (E, *Bc-GFP*), *domeMeso-GAL4,UAS-2xEYFP* (F, *domeMeso>GFP*). Cells were counterstained with phalloidin (A: green, B-F: red) and DAPI (blue). Scale bar 200µm. **(G)** Quantifications of the proportion of blood cells expressing the indicated transgenes in 5 or 35-day old flies. Means and standard deviations from at least 3 independent experiments are represented. **(H-P)** High magnification views showing the different types of blood cells observed in bleeds from 5-day old flies carrying the following combination of transgenes: *srpHemo-His2A-RFP, HmlΔ-GAL4,UAS-2xEYFP* (H), *srpHemo-His2A-RFP, BcF2-GFP* (I), *srpHemo-His2A-RFP, domeMeso-GAL4,UAS-2xEYFP* (J), *Pxn-RedStinger, HmlΔ-GAL4,UAS-2xEYFP* (K), *Pxn-RedStinger, BcF2-GFP* (L), *Pxn-RedStinger, domeMeso-GAL4,UAS-2xEYFP* (M), *HmlΔ-GAL4,UAS-2xEYFP, BcF6-mCherry* (N), *HmlΔ-GAL4,UAS-2xEYFP, crq-RedStinger* (O), *Cg25C-GFP, BcF6-mCherry* (P). Cells were counterstained with DAPI (blue) and phalloidin (green). Scale bar 20µm. **(Q)** Quantifications of the proportion of cells expressing the indicated combination of markers in 5 or 35-day old flies. **(R)** Schematic representation of the adult hematopoietic landscape. The proportion of cells expressing each marker is indicated.

Next, we tested whether lamellocytes can be retrieved in the adult. Intriguingly, we found that *msnF9* reporter, whose activity is induced in lamellocytes in the larvae (Tokusumi et al., 2009b), was expressed in ±90% of the adult hemocytes (Supplementary Figure 1G). However, we did not observe cells with high actin content and/or expressing the ß-integrin Myospheroid (Mys), two characteristic features of lamellocytes (Irving et al., 2005) (Supplementary Figure 2C, G), suggesting that lamellocytes are not present and that the *msnF9* reporter is constitutively active in adult blood cells. Since lamellocytes are absent in healthy larva but massively induced in response to infection by eggs of the parasitoid wasp *Leptopilina boulardi* (Lanot et al., 2001), we assessed their presence in adult flies that survived wasp infestation. However, ß-integrin and phalloidin staining did not reveal the presence of any lamellocyte in the adult, while they were readily recovered in bleeds from infected larvae (Supplementary Figure 2B, D, F and H). Thus, it appears that lamellocytes do not persist during adulthood.

Finally, to reveal the presence of putative blood cell progenitors, we made use of *tepIV-GAL4, Ance-GFP*, *dome^PG125^-GAL4* and *domeMeso-GAL4*, all of which have been shown to label larval prohemocytes in the lymph gland (Avet-Rochex et al., 2010; Cho et al., 2020; Krzemien et al., 2007; Louradour et al., 2017; Rodrigues et al., 2021) or in peripheral blood cells (Fu et al., 2020). Whereas we did not detected any expression of *tepIV-GAL4*, *Ance-GFP* or *dome^PG125^-GAL4* in adult hemocytes (Supplementary Figure 1H-J), we found that *domeMeso-GAL4* was expressed in ±4% of cells both in 5 and 35-day old flies (Figure 3F, G). All the *domeMeso^+^* cells exhibited a characteristic fusiform shape with one or two filopodial extensions and an elongated nucleus (Figure 3F, J’, J’’’, M’’, M’’’). Importantly, none of the fusiform cells expressed the blood cell differentiation markers tested above (*Pxn, Cg25C, Hml, crq, Bc, lz* and *msn*), but they expressed *srp*, albeit at lower level than the other blood cells (Figure 3H-P and Supplementary Figure 3A-H). Also, immunostaining against the plasmatocyte marker P1/NimC1 as well as *in situ* hybridization against *Pxn* or against the crystal cell marker *Bc/PPO1* confirmed that *domeMeso^+^*/fusiform cells did not express these blood cell differentiation markers (Supplementary Figure 3I-K), strongly suggesting that they could be blood cell progenitors.

In sum, our data indicate that the adult hematopoietic system is essentially composed of plasmatocytes (±85%) and crystal cells (±10%) and could harbor a small population of undifferentiated cells (±4%). Furthermore, adult blood cells can be classified in different subcategories (depending on the expression of *Pxn, Hml, crq, Bc* and *domeMeso*) whose proportions are relatively stable between young and old flies (Figure 3R).

### Abdominal hematopoietic hubs have a similar composition as adult bleeds

Since adult bleeds may not accurately reflect the composition of the adult hematopoietic system, we also assessed the expression of some of the above reporters in the dorsal abdomen. As shown in Figure 4A-G, *srp, Pxn, Cg25C, Hml, crq, Bc*, or *domeMeso* expression was observed as dispersed cells in the abdomen, especially in segment A1 to A3 along the heart, in so-called “hematopoietic hubs”. Srp expression was also prominent in pericardial cells (Figure 4A, arrows). In the hematopoietic hubs, *Hml^+^* cells represented 65% of the hemocytes as judged by co-labelling with *srp* or *Pxn* and no *Hml^+^ Pxn^-^* or *Hml^+^ srp^-^* cells were observed (Figure 4H, K and P). Crystal cells (*Bc*^+^) and putative prohemocytes (*domeMeso*^+^) composed ±10% and ±3% of the hemocytes respectively (Figure 4I, J and P). 90% of the crystal cells expressed *Cg25C* and 55% of them expressed *Hml* (Figure 4M, O and P). All the *domeMeso^+^* cells expressed *srp* at low levels and none expressed *Pxn* (Figure 4J, L and P). Similar observations were made when we analyzed the expression of these makers in 35-day old flies (Figure 4P). These results are consistent with our observations on adult bleeds and indicate that the adult hematopoietic system is composed of different subpopulations of plasmatocytes and crystal cells as well as a few putative blood cell progenitors.

**Figure 4.**
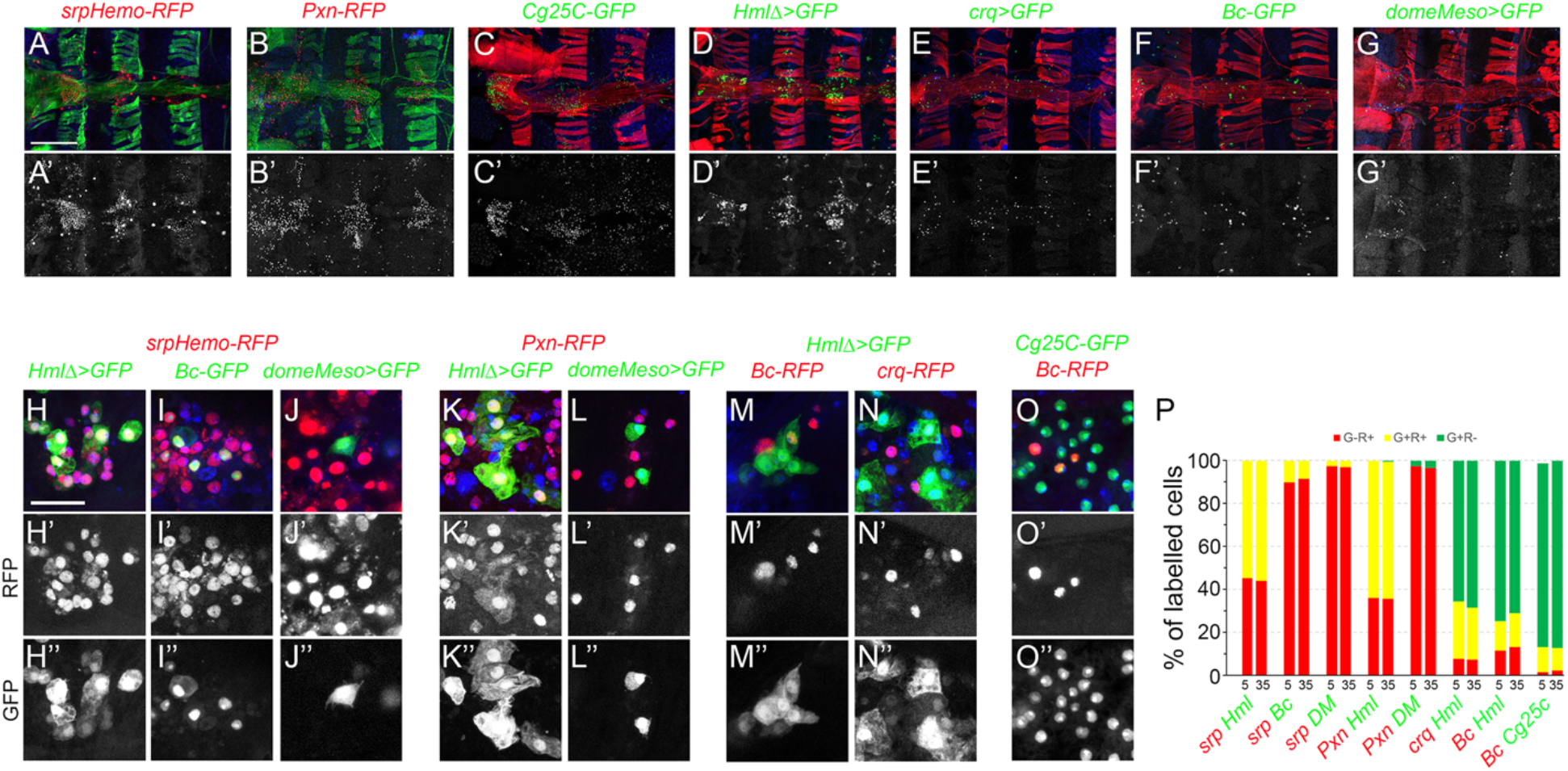
Analysis of blood cell transgenic markers in the adult abdomen. **(A-F)** Confocal views of the abdominal segments A1 to A4 of 5-day old females carrying the following transgenes: *srp-Hemo-His2A-RFP* (A), *Pxn-RedStinger* (B), *Cg25C-GFP* (C), *HmlΔ-GAL4,UAS-2xEYFP* (D), *crq-GAL4,UAS-2xEYFP* (E), *BcF2-GFP* (F), *domeMeso-GAL4,UAS-2xEYFP* (G). Cells were counterstained with phalloidin (A, B: green; C-G: red) and DAPI (blue). Arrows in (A) indicate some of the pericardial cells expressing *srp-Hemo*. The lower panels show only the transgene expression. Scale bar 200µm. **(H-O)** High magnification views of the hemocytes in the abdominal hematopoietic hubs (heart region in A1 or A2 segments) of 5-day old females carrying the following transgenes: *srpHemo-His2A-RFP, HmlΔ-GAL4,UAS-2xEYFP* (H), *srpHemo-His2A-RFP, BcF2-GFP* (I), *srpHemo-His2A-RFP, domeMeso-GAL4,UAS-2xEYFP* (J), *Pxn-RedStinger, HmlΔ-GAL4,UAS-2xEYFP* (K), *Pxn-RedStinger, domeMeso-GAL4,UAS-2xEYFP* (L), *HmlΔ-GAL4,UAS-2xEYFP, BcF6-mCherry* (M), *HmlΔ-GAL4,UAS-2xEYFP, crq-RedStinger* (N), *Cg25C-GFP, BcF6-mCherry* (O). The lower panels show only the red (H’-O’) or green (H”-O”) channel. Nuclei were stained with DAPI (blue). Scale bar 50µm. **(P)** Quantifications of the proportion of cells expressing the indicated combination of markers in 5 or 35-day old flies.

### Hml is expressed in a sub-population of adult hemocytes

The *Hml* driver has been used in several instance to manipulate gene expression in adult blood cells and/or test the function of these cells, notably by generating “hemoless” adults through the expression of apoptosis inducer in blood cells (Ayyaz et al., 2015; Charroux and Royet, 2009; Defaye et al., 2009; Ghosh et al., 2015; Sanchez Bosch et al., 2019; Woodcock et al., 2015). Yet our results suggest that this driver is not active in every adult hemocytes but rather labels a specific subpopulation. To confirm this hypothesis, we expressed the apoptotic inducer Bax under the control of *Hml* either during the whole development or only from day 1 of adulthood by using a *tubGAL-80^ts^* transgene to restrict its expression temporally, and we assessed the presence of hemocytes in 10-day old adult flies. *Hml*-driven expression of Bax specifically during adulthood caused an almost total ablation of the *Hml^+^* blood cells, but *Pxn^+^* cells were still present in adult bleeds and abdominal preparations (Figure 5B, D). When Bax was expressed from embryogenesis onward, we observed an efficient ablation of *Hml^+^* cells and very few *Pxn^+^* blood cells in adult abdominal hubs but, in contrast with the normal situation, we recovered many *Pxn^-^* cells when these individuals were bled (Figure 5F, H). While some of these *Pxn^-^* cells were fusiforms, as in the control, most had the typical round morphology of differentiated hemocytes, suggesting that the chronic expression of Bax in *Hml^+^* cells leads to the repression of *Pxn*. Consistent with this hypothesis, immunostaining against P1/NimC1 showed that many cells expressing this plasmatocyte marker were still present in adult bleeds and abdominal preparations following the constitutive expression of Bax in *Hml^+^* cells (Figure 5J, L and Supplementary Figure 4). These results show that killing *Hml^+^* blood cells is not sufficient to generate “hemoless” flies and might alter the gene expression program of the remaining blood cells. These data also indicate that during adulthood *Hml^+^Pxn^+^* and *Hml^-^Pxn^+^* cells form two distinct populations and that fusiform cells are not derived from *Hml^+^* cells.

**Figure 5.**
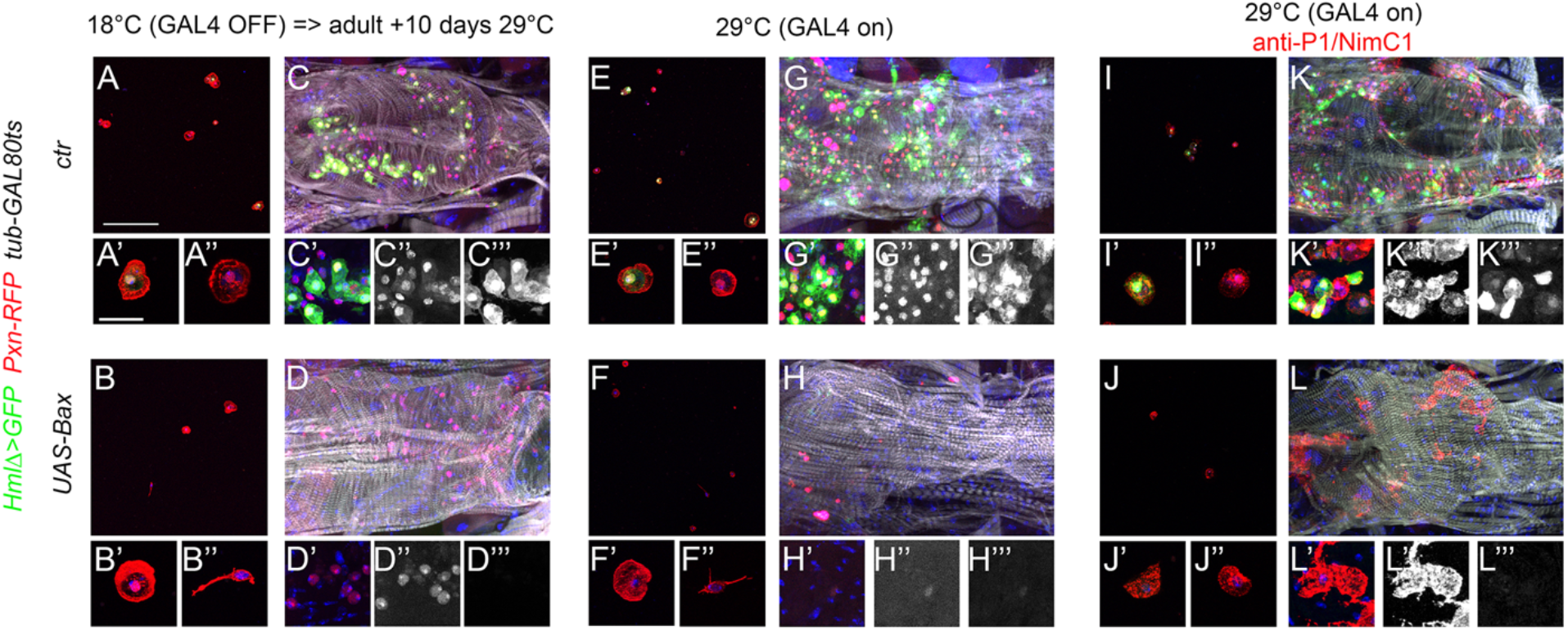
Adult blood cells are still present following the ablation of Hml^+^ hemocytes. Confocal views of adult bleeds and abdominal hubs from *tub-GAL80ts, Pxn-RedStinger, HmlΔ-GAL4,UAS-2xEYFP* (A, C, E, G, I, K) and *tub-GAL80ts, Pxn-RedStinger, HmlΔ-GAL4,UAS-2xEYFP, UAS-Bax* (B, D, F, H, J, L) flies expressing (or not) Bax in *Hml^+^* cells only during adulthood (A-D) or throughout their life (E-L). Cells were stained with phalloidin (A, B, E, F, I, J: red; C, D, G, H, K, L: white) and nuclei with DAPI (blue). (I-L): immunostaining against P1/NimC1 (red). Lower panels show high magnification views of the hemocytes. (C”, D”, G”, H”, K”, L”): red channel only. (C”’, D”’, G”’, H”’, K”’, L’”): green channel only. Upper panels: scale bar 200µm; lower panels: scale bar 20µm.

### *domeMeso^+^* “undifferentiated hemocytes” are not derived from larval blood cell progenitors but from the lymph gland posterior signaling center

A previous report showed that a small fraction of cells expressing Srp did not express the blood cell differentiation markers P1/NimC1 (for plasmatocytes) or Hindsight (for crystal cells), suggesting that they could be hematopoietic progenitors (Ghosh et al., 2015). In addition, these cells were proposed to derive from the posterior lobes of the larval lymph gland based on lineage tracing experiments using the *col-GAL4* driver *GMR13A11*. While *col-GAL4*-derived cells from the larval lymph gland were shown to contribute to adult plasmatocytes and crystal cells (Ghosh et al., 2015), it remains to be demonstrated that these cells can proliferate and/or differentiate during adulthood and thus constitute a genuine adult hematopoietic progenitor population. Our results above strongly suggested that fusiform*/domeMeso^+^* cells could correspond to this putative hematopoietic progenitor population. To test this hypothesis, we first assessed their origin. Using the G-TRACE cell lineage tracing technique (Evans et al., 2009) with either *GMR13A11* or the classically used *pcol85 col-GAL4* lines, we found that fusiform cells belonged to the *col* lineage-traced cells (Figure 6A, D), indicating that they correspond to the previously described adult blood cell progenitors. Interestingly we found that these cells maintain *col* driver expression, while we only observed past/lineage-traced expression in round hemocytes (Figure 6B, E). This result was confirmed by immunostaining, which revealed that Col was expressed in fusiform/*domeMeso^+^* cells, but not in other adult blood cells (Figure 6G, H and Supplementary Figure 5A, B). Furthermore, in accordance with the above characterization of *domeMeso^+^* cells, *col^+^* cells were *Pxn^-^* both in adult bleeds and in abdominal hematopoietic hubs (Figure 6I-K).

**Figure 6.**
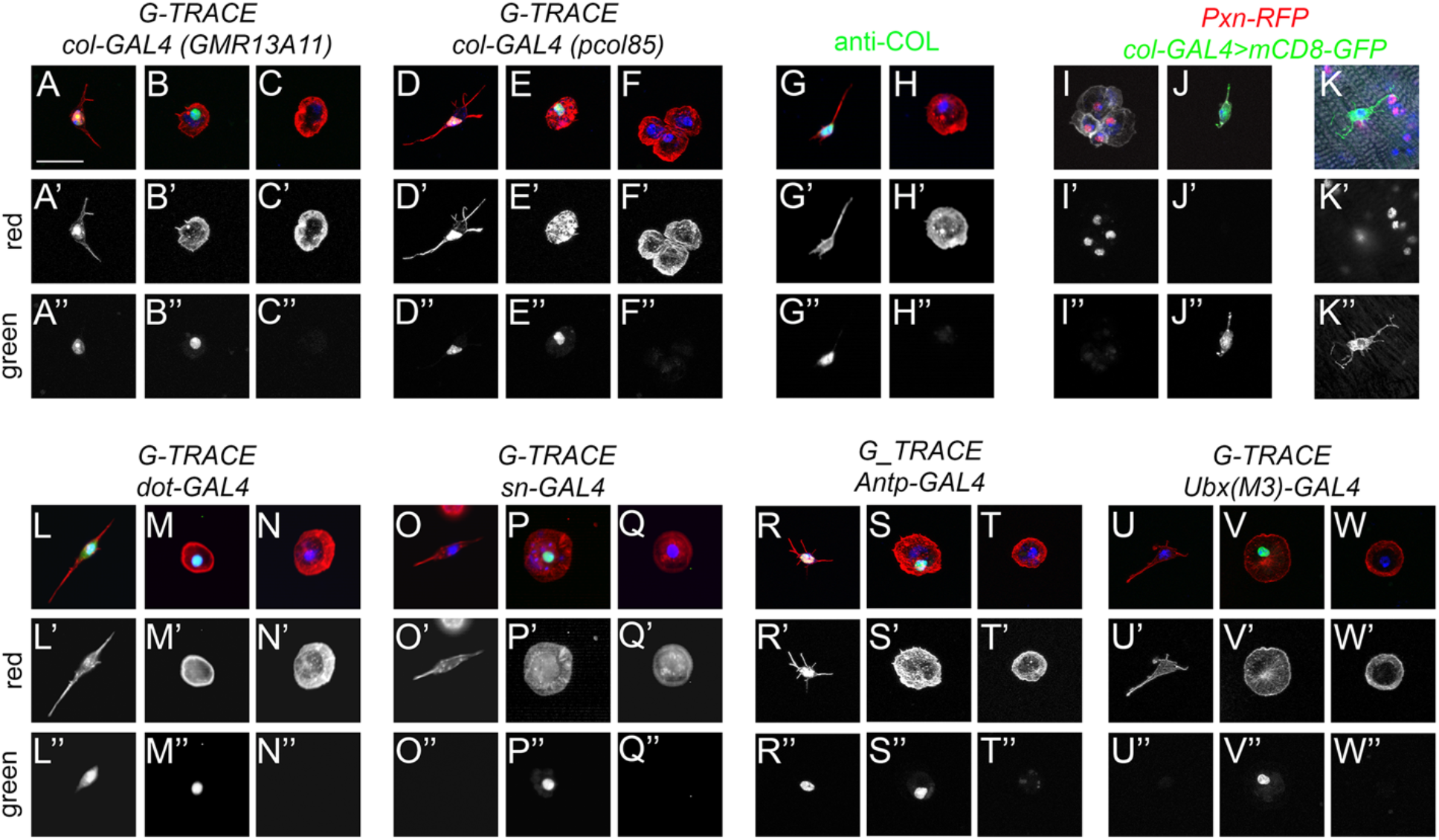
Fusiform cells are derived from the lymph gland posterior signaling center. **(A-F and L-W)** Cell lineage analyses of adult blood cells origin. Confocal views showing the different kinds of blood cells recovered in adult flies expressing the G-TRACE reporter transgenes under the control of the indicated drivers. Present and past expression of the driver are revealed by nuclear RFP (red, A’-F’ and L’-W’) and nuclear GFP (green, A”-F” and L”-W”) respectively. **(G-H)** Expression of Col (green) in adult blood cells as revealed by immunostaining. **(I-K)** Confocal views of adult bleeds (I, J) and abdominal hematopoietic hub (K) from *Pxn-RedStinger, pcol85-GAL4,UAS-mCD8-GFP* flies. (A-W): scale bar 20µm. Cells were counterstained with phalloidin (A-H and L-W: red; I-K: white) and DAPI (blue).

These results are thus consistent with the hypothesis that *col^+^* progenitors could give rise to differentiated hemocytes. Importantly though, the *col-GAL4* lines *pcol85* and *GMR13A11* are not only expressed in blood cell progenitors present in the lymph gland posterior lobes but also in the PSC (Benmimoun et al., 2012; Ghosh et al., 2015; Rodrigues et al., 2021). Moreover, recent single-cell RNA-seq analyses revealed that PSC-like cells expressing *col* and *Antp* are present in circulating larval blood cells (Cattenoz et al., 2021). Hence, these putative adult hematopoietic progenitors could have diverse origin. First, to test whether they are derived from the embryo/peripheral hemocytes or from the lymph gland, we performed lineage-tracing experiments with *sn-GAL4*, which is specifically expressed in embryo-derived hemocytes (Avet-Rochex et al., 2010), and *dot-GAL4*, which is specifically expressed in lymph gland-derived cells (Honti et al., 2010). Consistent with the idea that adult blood cells are derived from both waves of hematopoiesis (Holz et al., 2003; Sanchez Bosch et al., 2019), we observed lineage-traced expression in round hemocytes with both drivers (Figure 6M, P). In contrast, fusiform/Col^+^ cells were lineage-traced only with *dot-GAL4* (Figure 6L, O and Supplementary Figure 5C, D), demonstrating that they are solely derived from the lymph gland. Next, we asked whether fusiforms cells are derived from the lymph gland posterior lobes or the PSC using respectively *Ubx(M3)-GAL4*, which is expressed only in the posterior lobes (Rodrigues et al., 2021), and *Antp-GAL4*, which is expressed in the PSC but not in the posterior lobes (Evans et al., 2009; Rodrigues et al., 2021). Importantly, we found that fusiform cells were lineage-traced with *Antp-GAL4,* whereas *Ubx(M3)-GAL4* activity was traced only in round hemocytes (Figure 6R-W). Furthermore, immunostainings revealed that the PSC marker Antp is specifically expressed in *domeMeso^+^* blood cells (Supplementary Figure 5E, F). All together these results show that fusiform/*col^+^/domeMeso^+^* cells are derived from the PSC rather than from undifferentiated blood cells originating from the lymph gland posterior lobes.

### *domeMeso^+^* cells can differentiate and proliferate

Since *col* is required for PSC cell development (Crozatier et al., 2004), we tested its role for fusiform cell development. In *col^-/-^* adult flies, no *domeMeso^+^* cells (or fusiform cells) were present (Figure 7B). In addition, RNAi-mediated knock-down of *col* specifically in *domeMeso^+^* cells was sufficient to ablate this lineage (Figure 7C). To test whether the maintenance of *col* expression in *domeMeso^+^* cells is required in the adult, we used a *tub-GAL80^ts^* transgene to restrict *col* knock-down to adulthood. Following the inhibition of *col* expression for 48h in adult flies almost no *domeMeso^+^*/fusiform cells were retrieved, yet we observed a few cells that maintained *domeMeso* expression at lower levels and adopted a round shape similar to differentiated hemocytes (Figure 7F-I). Interestingly, in contrast with *domMeso^high^*/fusiform cells, these *domeMeso^low^* cells expressed the plasmatocyte marker P1/NimC1 (Figure 7D-I). These results show that *col* cell-autonomously controls *domeMeso^+^* cells development and suggest that its expression in these cells is continuously required to prevent their differentiation.

**Figure 7.**
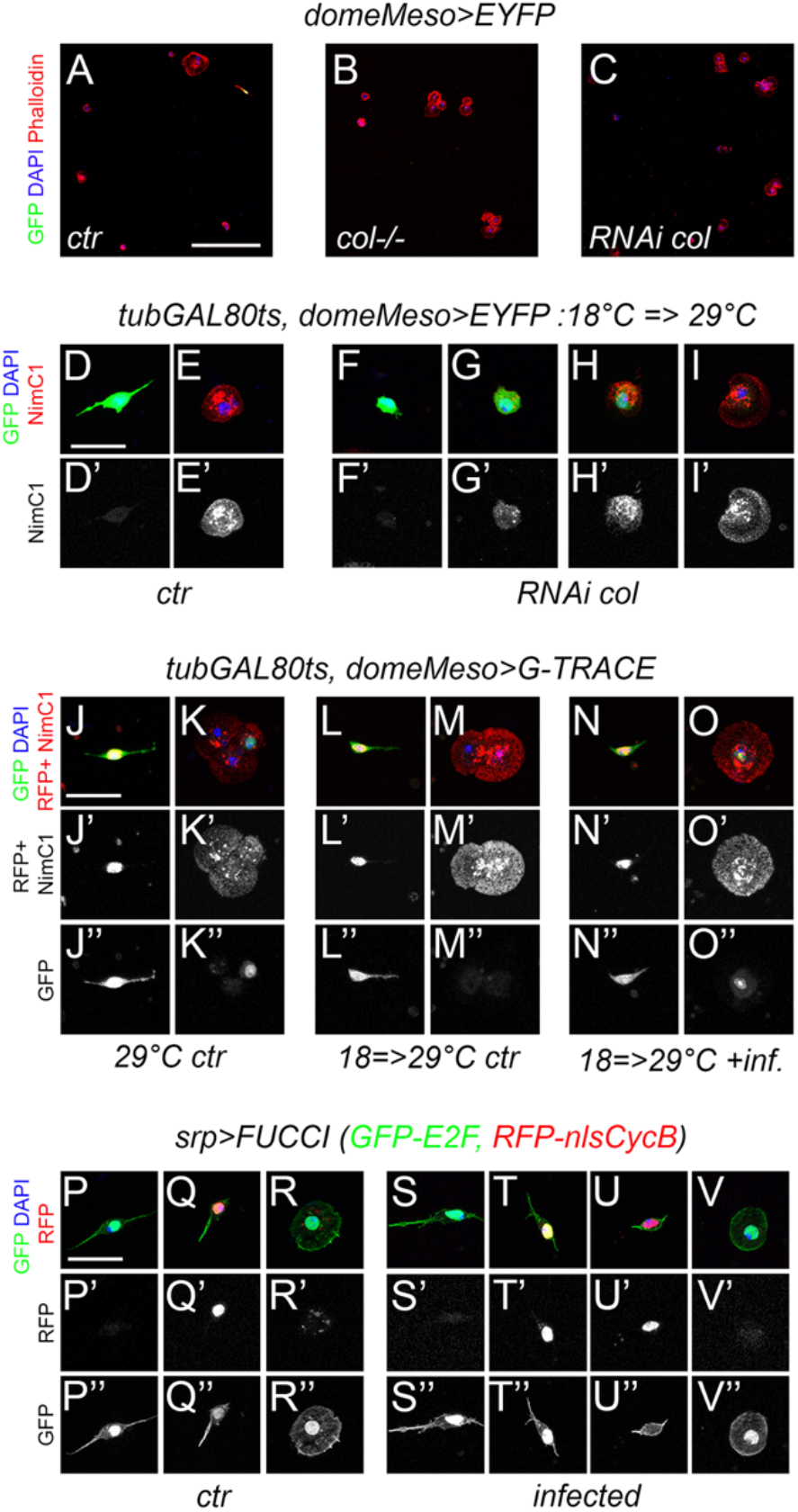
Fusiform cells can differentiate and proliferate. **(A-C)** Confocal views of blood cells from *domeMeso-GAL4, UAS-2xEYFP* (A, *ctr*), *col^-/-^, domeMeso-GAL4,UAS-2xEYFP* (B, *col^-/-^*) and *domeMeso-GAL4,UAS-2xEYFP, UAS-RNAi col* (C, *RNAi col*) adult flies. Blood cells were stained with phalloidin (red) and DAPI (blue). Scale bar 200µm. **(D-I)**. Blood cells from *tub-GAL80ts, domeMeso-GAL4,UAS2x-EYFP* adult flies expressing (D, E) or not (F-I) *col RNAi* only during adulthood. P1/NimC1 expression (red / lower panels) was revealed by immunostaining. **(J-O)** Blood cells from *tub-GAL80ts, domeMeso-GAL4, G-TRACE* flies raised at permissive temperature (J, K) or switched from restrictive to permissive temperature 24h after adult emergence (L, O). *domeMeso* live expression is revealed by nuclear RFP (red) and its lineage-traced expression by nuclear GFP (green). P1/NimC1 expression (red) was revealed by immunostaining. *ctr*: control non-infected flies, *inf*.: flies infected with *E. coli*. **(P-V)** Representative images of the cell cycle status observed in fusiform or round hemocytes from *srpD-GAL4,UAS-FUCCI* control (P-R) or *E. coli*-infected (S-V) adult flies. Cells in G1 phase express the nuclear GFP only, cells in G2/M express both nuclear GFP and RFP, cells in S phase express the nuclear RFP only. Cell morphology was visualized by phalloidin staining (green). (D-V) Nuclei were stained with DAPI. Scale bar 20µm.

In light of the above results, we used the G-TRACE system to test whether *domeMeso^+^* cells naturally differentiate during adulthood. Because *domeMeso-GAL4* is expressed in blood cells progenitors in the larval lymph gland (Louradour et al., 2017; Rodrigues et al., 2021), we used a *tubGAL80^ts^* transgene to temporally restrict its activity. When *domeMeso-GAL4* activity was left unrestricted (*i.e.* when flies were raised at 29°C from embryonic to adult stages), we observed lineage-traced expression in round cells (which expressed the plasmatocyte marker P1/NimC1) as well as live (and traced) expression in fusiform cells (Figure 7J, K). When *dome-GAL4* activity was restricted to the adult (*i.e.* when flies were raised at 18°C until adulthood and then switched at 29°C for up to two weeks), *domeMeso>G-TRACE* expression was observed in fusiform cells but not in other hemocytes (Figure 7L, M). Strikingly though, we found that upon infection of adult flies by *E. coli*, some round/NimC1^+^ cells were lineage-traced with *domeMeso* (Figure 7N, O). Thus, while *domeMeso^+^* cells do not differentiate at detectable rate during adulthood under normal conditions, they can give rise to plasmatocyte in response to an immune challenge.

Finally, it was proposed that infection of adult flies with *E. coli* can induce *Hml^+^* cells proliferation at low frequency (Ghosh et al., 2015), but these results were rebuked by Sanchez Bosch et al., who brought several lines of evidence that *Hml^+^* cells do not proliferate even in response to infection (Sanchez Bosch et al., 2019). Yet, *Hml* is not expressed in all the blood cells and notably not in *domeMeso^+^*/fusiform cells. We thus reassessed the cell cycle status of adult blood cells using *srpD-GAL4,* which is expressed in all the hemocytes, and the two-color FUCCI cell cycle indicator (Zielke et al., 2014). We found that round hemocytes were in G1 phase, while 70% of the fusiform cells were in G1/S and 30% in G2/M (Figure 7P-R). No blood cells were found to be in S phase, indicating that they do not proliferate in uninfected adults. Following infection with *E. coli*, round cells remained in G1 and the proportion of fusiform cells in G1 (68%) or G2/M (30%) was barely affected (Figure 7S-V). However, we repeatedly found a few fusiform cells in S phase (2%) (Figure 7U). These results thus indicate that fusiform cells can proliferate, albeit at low frequency, in response to an immune challenge.

## Discussion

In mammals, the life-long production of the different blood cell types relies on the presence of long-term hematopoietic stem and progenitor cells, which are specified in the embryo and reside in specific niches from where they can be mobilized to respond to the needs of the organism (Cool and Forsberg, 2019). While several aspects of hematopoiesis have been conserved from mammals to Drosophila (Banerjee et al., 2019), the presence of a similar long-term blood cell progenitor population in Drosophila remains elusive and the overall composition of the adult hematopoietic system is still a matter of debate. In particular, two successive publications came to different conclusions concerning the presence of hematopoietic progenitors and the mere existence of hematopoiesis in adult flies (Ghosh et al., 2015; Sanchez Bosch et al., 2019). Overall, the data we obtained indicate that the Drosophila adult hematopoietic system is essentially composed of different subpopulations of mature blood cells, including PSC-derived cells which seem to have some limited hematopoietic capacity, and concur with the hypothesis that no significant hematopoiesis takes place during adulthood.

Our results show that the bleeding protocol that we used ensures a swift and representative collection of the blood cell types present in the adult with minimal contamination from other tissues. This simple protocol is thus suitable to define the adult hemocyte gene repertoire without applying tissue dissociation and cell-purification protocols, which might alter gene expression. Thereby, we could show that adult hemocytes share an important part of their transcriptome with their larval parents, but also exhibit some clearly distinct features. Notably several blood cell markers are differentially expressed between adult and larval hemocytes. For instance, *NimC4* and *drpr*, which work together for apoptotic cell clearance in the embryo (Kurant et al., 2008), or *Fer1HCH* and *Fer2LHC*, which encodes the two components of the iron transporter ferritin and control larval blood cell differentiation (Yoon et al., 2017), are strongly up-regulated in adult hemocytes. Moreover, our findings highlight an unexpected degree of similarities between adult and embryonic blood cells as compared to larval blood cells. Indeed, the comparison of our results with the recently described embryo *versus* larval blood cell gene repertoire (Cattenoz et al., 2020) revealed that almost half of the genes differentially expressed in adult *versus* larval hemocytes are similarly regulated in embryonic *versus* larval blood cells. This is particularly striking for adult up-regulated genes as 422 (out of 864) of them are also more strongly expressed in embryonic hemocytes, indicating that adult blood cells reactivate a large number of embryonic blood cell markers. Among those, many are involved in energy metabolism. Actually embryo-derived hemocytes undergo a shift from glycolysis during embryogenesis to lipid ß-oxidation during the larval stages (Cattenoz et al., 2020) and our results strongly argue that a second metabolic shift occurs during adulthood. The important role of energy metabolism regulation in Drosophila blood cells is underscored by recent publications showing that fatty acid ß-oxidation is required for the differentiation of blood cell progenitors in the larval lymph gland (Tiwari et al., 2020) and that infection leads to changes in blood cell metabolism which are important for an effective immune response (Bajgar et al., 2015; Krejcova et al., 2019). Prominent changes in energy metabolism have been observed also in mammalian blood cells in response to development, aging, infection or cancer (Faas and de Vos, 2020; Nakamura-Ishizu et al., 2020; Rashkovan and Ferrando, 2019). Getting further insights into the pathways controlling these shifts will thus be of broad interest. More generally, the differences between embryonic, larval and adult blood cell gene expression programs likely reflect distinct biological functions which warrant further investigation.

Our analysis of different blood cell differentiation markers indicate that the adult hematopoietic system comprises different subtypes of plasmatocytes as well as ±10% of crystal cells, but no lamellocytes. These finding are consistent with previous reports (Ghosh et al., 2015; Kurucz et al., 2007; Lanot et al., 2001; Sanchez Bosch et al., 2019), and underscore the heterogeneity of the plasmatocyte/crystal cell populations, which can be subdivided according to the expression of *Hml*, *crq* and, to a lesser extent, *Cg25C*. It will be interesting to analyze whether these subpopulations have distinct functions for instance in the immune response or the removal of apoptotic cells, as recently shown for embryonic macrophage subpopulations (Coates et al., 2021). The proportion of these subpopulations was similar between young and aged flies, suggesting that they do not represent differentiation intermediates inherited from pupal hemocytes. Along that line, our cell ablation experiments argue against the idea that all the adult hemocytes are derived from *Hml^+^* cells or that adult *Hml^+^Pxn^+^* cells differentiate into *Hml^-^/Pxn^+^* cells during adulthood. Besides, our observations are in line with previous reports showing that adult blood cells do not proliferate (Lanot et al., 2001; Sanchez Bosch et al., 2019) and that their number decreases with aging (Mackenzie et al., 2011; Sanchez Bosch et al., 2019; Woodcock et al., 2015). Therefore, it seems that the adult hematopoietic system is essentially composed of mature blood cells.

Ghosh *et al*. observed some Srp^+^ blood cells that did not express the differentiation markers NimC1 or Hnt and were lineage-traced with *col-GAL4* (Ghosh et al., 2015). Although the differentiation and proliferative potentials of these cells during adulthood were not assessed, the authors concluded that blood cell progenitors derived from the lymph gland posterior lobes persist during adulthood. Our results contradict this model. Indeed, while we also observed the presence of a small proportion of *srp^+^* cells that do not express hematopoietic differentiation marker and are lineage-traced as originating from the lymph gland, we demonstrated that these cells are actually derived from the PSC rather than from the posterior lobes. Actually, they exhibit a characteristic fusiform shape with philopodial extensions that is reminiscent of PSC cells (Krzemien et al., 2007; Mandal et al., 2007), and they express Col, which is required for PSC development but also for their maintenance in the adult. Moreover, although these cells express *domeMeso*, they do not express other prohemocytes markers (*tep4*, *Ance* or *dome-GAL4),* and we did not find evidence that they differentiate during adulthood in normal conditions. Nonetheless these cells hold some hematopoietic potential. Notably, upon *E. coli* infection, some of them can enter S phase and they can differentiate into plasmatocyte as evidenced by NimC1 induction and morphological changes. Along the same line, infection of larvae with *E. coli* was shown to induce PSC cells proliferation and to affect their morphology (Khadilkar et al., 2017). However, given the low number of PSC-derived cells and the limited response we observed in infected adults, it is unlikely that *de novo* production of mature blood cells is their main function. In the lymph gland, the PSC forms a highly specialized cluster of cells that do not contribute to the general pool of blood cells but control their fate, in particular in response to immune challenges (Benmimoun et al., 2015; Crozatier et al., 2004). It is tempting to speculate that adult PSC-derived cells may have similar functions in the adult.

In sum, our results strongly suggest that the Drosophila adult hematopoietic system does not harbor a true blood cell progenitor population with significant proliferation and differentiation potential. Still, different types of mature, potentially specialized, blood cells are present in the adult. Along that line it will be interesting to decipher the function of the PSC-derived population, as these cells have been shown to control the cellular immune response to some specific immune challenges in the larva and may play a similar role in the adult. Finally, it is expected that applying single cell RNA-sequencing approaches to the adult hematopoietic system will bring a better description of its constituents and could reveal some of its functional features.

## Methods

### Fly strains

The following strains were used in this study: *w^1118^* (BL3605), *srpD-GAL4* (Crozatier et al., 2004; Waltzer et al., 2002), *srpHemo-H2A-3xmCherry* (Gyoergy et al., 2018) (BL78361), *HmlΔ-GAL4* (Sinenko and Mathey-Prevot, 2004) (BL30141 and BL30140), *crq-GAL4* (Olofsson and Page, 2005) (BL25041), *lz-GAL4* (Lebestky et al., 2000) (BL6313), *sn-GAL4* (Avet-Rochex et al., 2010; Zanet et al., 2012), *dot-GAL4* (Kimbrell et al., 2002) (BL6902), *dome^PG125^-GAL4* (Bourbon et al., 2002), *domeMeso-GAL4* (Oyallon et al., 2016), *tep4-GAL4* (Avet-Rochex et al., 2010) (DGRC #105442), *Ance^MiMiC^-GFP* (BL59829), *pcol85-GAL4* (Crozatier et al., 2004), *col(GMR13A11)-GAL4* (BL49248), *Antp-GAL4* (Mandal et al., 2007), *Ubx(M3)-GAL4* (de Navas et al., 2006), *Pxn-GAL4* (Stramer et al., 2005), *Pxn-RedStinger* (Rodrigues et al., 2021), *BcF2-GFP* (Gajewski et al., 2007), *BcF6-mCherry* (Tokusumi et al., 2009a), *msnF9-mCherry, msnF9-GFP* (Tokusumi et al., 2009b), *Cg-GAL4* (Asha et al., 2003) (BL7011), *Cg25C-GFP* (Sorrentino et al., 2007)*, UAS-RNAi col* (Baumgardt et al., 2007), *kn^col1^* and *kn^col1^P{col5-cDNA}* (Crozatier and Vincent, 1999), *UAS-Bax* (Gaumer et al., 2000), *UAS-2xEGFP* (BL6874), *UAS-RedStinger* (BL8546), *UASm-mCD8GFP* (BL5138), *tubP-Gal80^ts^* (BL7018)*, G-TRACE* (*UAS-RedStinger, UAS-FLP, Ubi-p63E(FRT.STOP)Stinger)* (BL28281), *UAS-FUCCI (UAS-GFP.E2f1.1-230, UAS-mRFP1.NLS.CycB.1-266)* (BL55110).

Drosophila stocks and crosses were maintained on standard fly medium (75 g/l organic corn flour, 28 g/l dry yeast, 40g/l sucrose, 8 g/l agar, 10ml/l Moldex 20%). All crosses and collections were performed at 25°C with the exception of experiments involving *tub-GAL80^ts^*, which were performed at 18°C before transferring the progenies to 29°C as indicated in the results section.

### Adult blood cells preparations

To bleed adult flies, aged-matched individuals were anesthetized, washed in 70% ethanol and air dried before cutting the last abdominal segment with a clean scalpel. Then a fine glass needle was inserted in the anterior part of the thorax and PBS was perfused under air pressure. The flushed hemocytes were collected in a 12-well plate (Nunc) containing 600µl PBS and a round glass coverslip. In general, 6 drops of PBS (±60µl) were collected per fly and 5 flies were bled in a single well. The plates were centrifuged at 1000 rpm for 2 min before adding 300µl of 16% formaldehyde for 30 min. For *in situ* observation of the blood cells in adult abdomen, the flies were dissected essentially as described in (Ghosh et al., 2015). Briefly: flies were anesthetized, stuck on their dorsal side in paraffin-coated plastic dish and dissected in PBS. The wings and most of the thorax were discarded and the ventral abdomen was incised on both sides with thin scissors before delicately removing the gonads and the gut. The dissected abdomens were fixed in 4% formaldehyde for 30 min. Samples were then processed as described below for immunofluorescence and/or confocal imaging.

### Immunostainings and *in situ* hybridizations

Immunostaining and RNA *in situ* hybridizations were essentially performed as described in (Miller et al., 2017). The following antibodies were used: mouse anti-P1/NimC1 (Kurucz et al., 2007), mouse anti-Col (Krzemien et al., 2007), mouse anti-ß integrin/Myospheroid (CF6.G11, DSHB), mouse anti-Antp (8C11, DSHB), rabbit anti-GFP (Torrey Pines), anti-DIG coupled to alkaline phosphatase (Roche) and Alexa-fluor labelled secondary antibodies (Molecular Probes). Nuclei were stained with DAPI and the actin cytoskeleton with fluorescently labeled phalloidin (Molecular Probes). The slides were mounted in Vectashield (Vector) and observed under a confocal microscope (Zeiss LSM800 or Leica SP5). Confocal images are displayed as maximum intensity projection of Z-stacks.

### Bacterial and wasp infections

*E. coli* (DH5*α* strain) were grown overnight in LB broth and pelleted at 4000 rpm for 15 min. A fine tungsten needle was dipped in the bacterial pellet and inserted into the dorsal abdomen or under the wing hinge of anesthetized 1-week old females. A heat-sterilized tungsten needle was used to prick the flies that were used as non-infected control. Three groups of 10 flies were used in each experiment. The flies were then reared for 16h (FUCCI experiments) to 96h (G-TRACE experiments) and dissected or bleed as described above. For wasp infestations, late second instar larvae were subjected to parasitism by ±10 *Leptopilina boulardi* (strain G486) females for 2h as described in (Benmimoun et al., 2015). Successful infestation was assessed by looking for melanotic nodule 48h later. The corresponding larvae were transferred to fresh vial and the surviving adult flies were bled 5 days after emergence.

### RNA-sequencing

For RNA-seq experiments, wandering third instar larvae or 4 to 5-day old adult *w^1118^* females were used to prepare independent biological triplicates. Adult blood cells were retrieved by perfusing the flies with PBS as described above and directly collected in a 1.5ml Eppendorf tube on ice. Peripheral larval blood cells were collected by delicately peeling the dorsal cuticle with forceps and dripping the larva in a 5µl drop of PBS on parafilm. Upon microscopic examination, we found that >90% of the cells retrieved with this protocol are hemocytes (*Pxn-RedStinger^+^*), with no visible contamination by fat body cells. The anterior lobes of the larval lymph glands were dissected in PBS. Blood cells and lymph glands were transferred to 1.5ml Eppendorf tubes, pelleted by centrifugation at 1000 rpm for 2min and processed for RNA extraction using Qiagen RNeasy Plus Micro kit (#74034). For each replicate, ±100 adult flies and 20 larvae were used. RNA samples were run on Agilent Bioanalyzer to assess sample quality and concentration. Samples were converted to cDNA using Nugen’s Ovation RNA-Seq System (Catalogue # 7102-A01). Libraries were generated using Kapa Biosystems library preparation kit (#KK8201) and multiplexed libraries were sequenced on a 1×75 flow cell on the HiSeq2000 platform (Illumina). Reads were filtered and trimmed to remove adapter-derived or low-quality bases using Trimmomatic and checked again with FASTQC. Illumina reads were aligned to Drosophila reference genome (dm6 Ensembl release 70) with Hisat2. Read counts were generated for each annotated gene using HTSeq-Count. RPKM (Reads Per Kilobase of exon per Megabase of library size) values were calculated using Cufflinks. Reads normalization, variance estimation and pair-wise differential expression analysis with multiple testing correction was conducted using the R Bioconductor DESeq2 package. Heatmaps and hierarchical clustering were generated with “pheatmap” R package. Gene ontology enrichment analyses were performed using Genomatix. The RNA-seq data were deposited on GEO under the accession number GSE174107.

### Real-time quantitative PCR

For RT-qPCR, RNA samples were prepared from adult or larval bleeds and larval lymph glands (dissected as described above) using RNeasy kit (Qiagen) with an additional on-column DNAse treatment with RNase-Free DNase Set (Qiagen). Reverse transcription was performed with SuperScript IV Reverse Transcriptase (ThermoFisher) according to manufacturers instruction on 100ng of RNA and using a mix (1:1) of random primers (Invitrogen) and oligo dT (Promega). qPCRs were performed with SsoFast EvaGreen (Biorad) on a LightCycler 480 Instrument II (Roche Life Science). The sequences of the primers used to assess the expression of the different genes are provided in Supplementary Table 7. qPCR data were analyzed with double delta Ct method and gene expressions were normalized to *rp49 (*synonym*: RpL32)*. All experiments were performed using biological triplicates.

## Acknowledgments

We thank Toulouse RIO and Clermont-Ferrand CLIC platforms for assistance with confocal imaging. We thank all our former colleagues for valuable discussions and the Drosophila community, the Drosophila Bloomigton Stock Center and the Developmental Study Hybridoma Bank (DSHB) for fly stocks and antibodies. This work was supported by grants from the Agence Nationale de la Recherche, i-Site CAP20-25 and the Indo-French Centre for the Promotion of Advanced Research to LW. MB was supported by fellowships from the Université Clermont-Auvergne and the Fondation pour la Recherche Médicale (FRM).

