## Supplementary figures for "Characterization of the Drosophila adult hematopoietic system reveals a rare cell population with differentiation and proliferation potential"

### Supplementary files

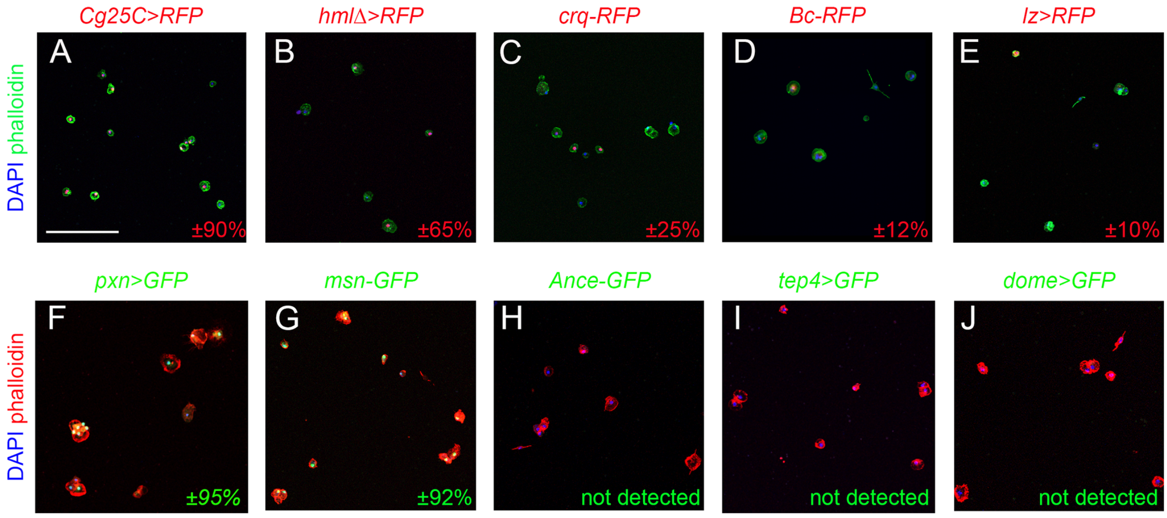


**Supplementary Figure 1.** Confocal views showing representative bleeds from 5-day old flies carrying the following transgenes: *Cg-GAL4,UAS-RedStinger* (A, *Cg25C>RFP*), *HmlΔ-GAL4,UAS-RedStinger* (B, *HmlΔ>RFP*), *crq-RFP* (C), *BcF6-mCherry* (D), *lz-GAL4,UAS-RedStinger* (E, *lz>RFP*), *pxn-GAL4,UAS-2xEYFP* (F, *pxn>GFP*), *msnF9-GFP* (G, *msn-GFP*), *Ance-GFP* (H), *tep4-GAL4,UAS-2xEYFP* (I, *tep4>GFP*), *dome*$^{PG125}$*-GAL4,UAS-2xEYFP* (J, *dome>GFP*). Cells were counterstained with phalloidin (A-E: red, F-J: green) and DAPI (blue). Scale bar 200µm. The percentage of cells expressing each transgene is indicated in the bottom right hand corner.

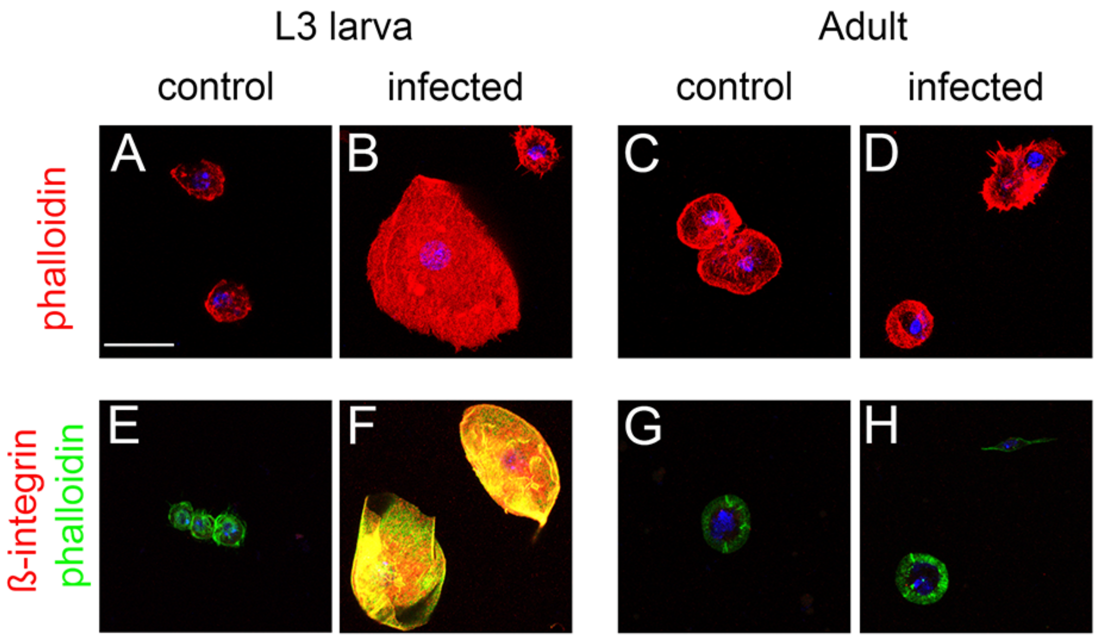


**Supplementary Figure 2**. Confocal views of larval (A, B, E, F) and adult (C, D, G, H) blood cells from control (A, E, C, G) or *L. boulardi*-infected (B, D, F, H) flies. (A-D) Blood cells morphology was revealed with phalloidin staining (red). (E-H) Blood cells were stained with phalloidin (green) and anti-ß-integrin (red). (A-H) Nuclei were stained with DAPI. Scale bar 20µm.

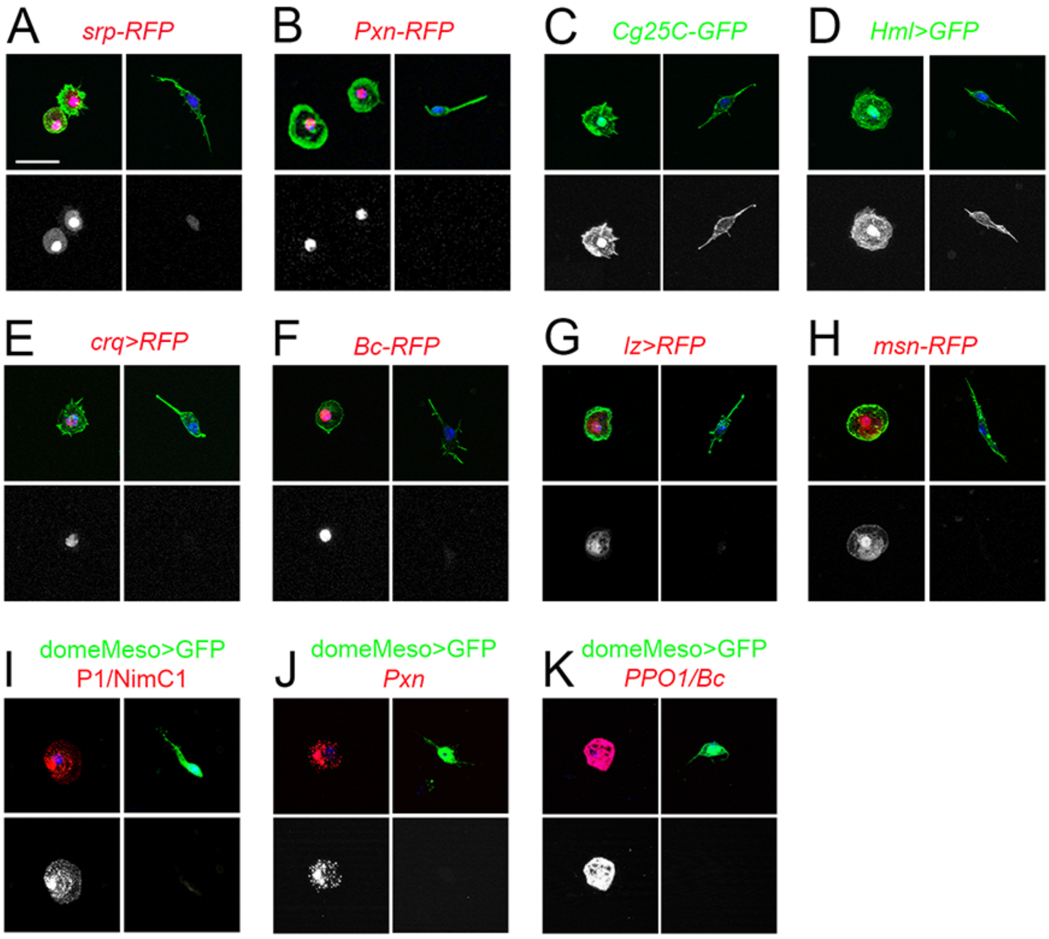


**Supplementary Figure 3**. (**A-H**) Confocal views of hemocytes from adult flies carrying the following transgenes: *srpHemo-His2A-RFP* (A), *Pxn-RedStinger* (B), *Cg25C-GFP* (C), *HmlΔ-GAL4,UAS2xEYFP* (D), *crq-GAL4,UAS-RedStinger* (E), *BcF6-mCherry* (F), *lz-GAL4,UAS-RedStinger* (G), *msnF9-mCherry* (H). Cells were counterstained with phalloidin (green) and DAPI (blue). Lower panels display the red (A, B, E-H) or green (C, D) channel only. **(I-K)** Confocal views of hemocytes from *domeMeso-GAL4,UAS-2xEYFP* adult flies following immunostaining against P1/NimC1 (I, red) or *in situ* hybridization against *Pxn* (J, red) or *PPO1/Bc* (K, red). Cell nuclei were stained with DAPI (blue). The lower panels display the red channel only. (A-K): scale bar 20µm. Left panels display round hemocytes and right panels show fusiform/*domeMeso*$^+$ cells.

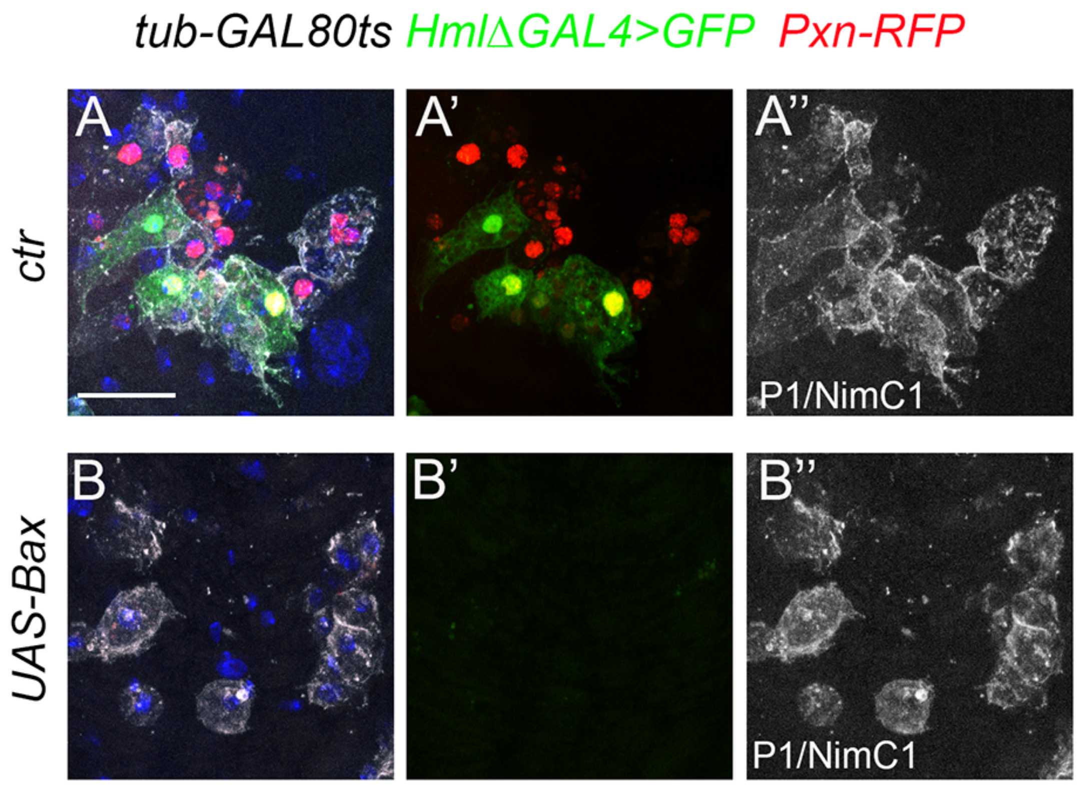


**Supplementary Figure 4.** Confocal views of abdominal hematopoietic hubs from *tub-GAL80ts, Pxn-RFP, HmlΔ-GAL4,UAS-2xEYFP* (A) and , *tub-GAL80ts, Pxn-RFP, HmlΔ-GAL4,UAS-2xEYFP, UAS-Bax* (B) flies raised at 29°C. Cells were stained with anti- P1/NimC1 (white) and DAPI (blue). (A', B'): green (GFP) and red (RFP) channels. (A", B"): white channel (P1/NimC1). Scale bar 20μm.

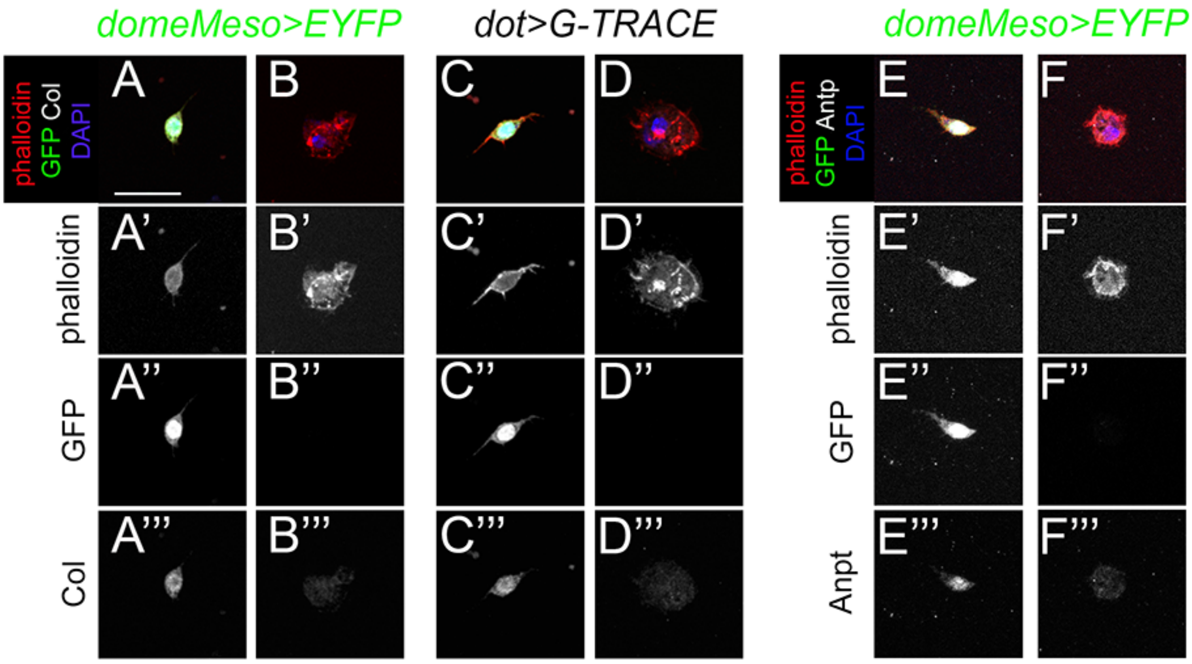


**Supplementary Figure 5.** Confocal views of Col (A-D) or Antp (E, F) expression in hemocytes from *domeMeso-GAL4, UAS-2xEYFP* (A, B, E, F) or *dot-GAL4,G-TRACE* (C, D) adult flies. (C, D) G-traced (past) activity of *dot-GAL4* is visualized by nuclear GFP expression. No live expression (nuclear RFP) was observed. Cells were stained with DAPI (blue) and phalloidin (red). Scale bar: 20μm.

**Supplementary Table 1**. List of genes expressed in adult hemocytes (RPKM>1 in all three samples).

**Supplementary Table 2.** List of genes expressed in larval peripheral hemocytes (RPKM>1 in all three samples).

**Supplementary Table 3.** List of genes expressed in larval lymph glands (RPKM>1 in all three samples).

**Supplementary Table 4.** List of hemocytes markers on Flybase (*expression*: hemocytes, plasmatocytes, crystal cells, lamellocytes or prohemocytes).

**Supplementary Table 5.** List of differentially expressed genes between adult, larval peripheral and lymph gland blood cells (fold change >2, adjusted p-value<0.01).

**Supplementary Table 6.** List of differentially expressed genes between embryonic and larval peripheral hemocytes (fold change >2, adjusted p-value<0.01) (from EMBL-EBI data set: E-MTAB-8702).

**Supplementary Table 7.** List of primers used for qPCR.
